# Robustness and timing of cellular differentiation through population-based symmetry breaking

**DOI:** 10.1101/578898

**Authors:** Angel Stanoev, Christian Schröter, Aneta Koseska

## Abstract

During mammalian development, cell types expressing mutually exclusive genetic markers are differentiated from a multilineage primed state. These observations have invoked single-cell multistability view as the dynamical basis of differentiation. However, the robust regulative nature of mammalian development is not captured therein. Considering the well-established role of cell-cell communication in this process, we propose a fundamentally different dynamical treatment in which cellular identities emerge and are maintained on population level, as a novel unique solution of the coupled system. Subcritical system’s organization here enables symmetry-breaking to be triggered by cell number increase in a timed, self-organized manner. Robust cell type proportions are thereby an inherent feature of the resulting inhomogeneous solution. This framework is generic, as exemplified for early embryogenesis and neurogenesis cases. Distinct from mechanisms that rely on pre-existing asymmetries, we thus demonstrate that robustness and accuracy necessarily emerge from the cooperative behaviour of growing cell populations during development.

## Main

Functional diversification during mammalian development arises through symmetry breaking events that characterize a transition from an initially homogeneous group of multilineage primed cells towards a heterogeneous population of differentiated cellular identities (Zhang and Hiiragi, 2018; Simon et al., 2018). These events are typically generated and the corresponding states are maintained through self-organized cooperative processes, whose respective dynamics cannot be deduced from the dynamical features of the individual cells (Zhang and Hiiragi, 2018; Kauffman, 1993). Even more, the relatively low information content of a system residing in a symmetrical homogeneous state implies that information originates with the generated cell types at the symmetry breaking event. The onset of this event must be accurately timed, where not only the specification of distinct cell fates is suitably determined, but also the number distribution of each type must be robustly established.

The observations that expression of mutually exclusive genetic markers distinguishes the differentiated fates among each other and from the multilineage primed state have however promoted the hypothesis that multistability on the level of single cells sets the dynamical basis for differentiation (Kauffman, 1969; Andrecut et al., 2011; Wang et al., 2011; Enver et al., 2009). The most common functional motif that accounts for bistability on a single cell level is a two-component genetic toggle-switch (Thomas, 1981; Cherry and Adler, 2000; Snoussi, 1998), whereas addition of self-activating loops (Huang et al., 2007; Bessonnard et al., 2014; Jia et al., 2017) gives rise to a third stable state – the multilineage primed co-expression state. Such single cell multistable circuits have been used to describe the Gata1/PU.1 switch that governs the lineage commitment in multipotent progenitor cells (Huang et al., 2007; Graf and Enver, 2009), the Cdx2/Oct4 switch in the differentiation of the totipotent embryo (Niwa et al., 2005), the T-bet/Gata3 switch in the specification of the T-helper cells (Huang, 2013), as well as the Gata6/Nanog switch in the branching process of inner cell mass (ICM) (Bessonnard et al., 2014; Chickarmane and Peterson, 2008). In these systems, pre-existing asymmetries, typically attributed to stochastic events or cell-to-cell heterogeneities, are assumed to be amplified by intercellular signaling, and are necessary to drive the individual cellular states out of the multilineage primed- and into one of the differentiated attractors (De Mot et al., 2016; De Caluwé et al., 2019). However, as all of the differentiated states are initially present in this description, symmetry breaking does not formally occur. Even more, experimental evidence suggests that cell-cell communication and signaling is crucial to obtain differentiated cell types during early mammalian development (Nichols et al., 2009; Yamanaka et al., 2010). Thus, cells do not function as isolated entities that process the information from the environment in a unidirectional input-output fashion, but operate as a joint dynamical system in which they continuously communicate by secreting growth factors or other signaling molecules. The principle of an emergent symmetry breaking mechanism that characterizes how cells differentiate into heterogeneous types while simultaneously accounting for the beginning and robustness of the process, is therefore still unclear.

We propose a dynamical mechanism where a population of identical cells breaks the symmetry due to cell-cell communication, giving rise to a novel heterogeneous dynamical solution that is different than the solutions of the isolated cells. We identified that the transition from homogeneous to a heterogeneous population is uniquely governed by a subcritical pitchfork bifurcation, resulting in the formation of an inhomogeneous steady state (IHSS) that reflects the cooperatively occupied differentiated cell fates. The formation of this new, population-based heterogeneous attractor additionally demonstrates how information is generated at a symmetry breaking event. Parameter organization in the vicinity of its bifurcation point enables cell number increase to trigger the symmetry breaking event in a self-organized manner, which renders timing of cellular differentiation – an emergent property of growing populations. Moreover, reliable cell proportions in the differentiated fates are an inherent feature of this symmetry-breaking solution. The proposed mechanism is generic and applies to systems with diverse gene expression dynamics in single cells, as we additionally demonstrate for a bistable circuit to describe the differentiation into epiblast (Epi) and primitive endoderm (PrE) states from the homogenous ICM during the blastocyst development stage of the mammalian preimplantation embryo, as well as for single cell oscillations, capturing the vertebrate neurogenesis dynamics.

## Results

### Heterogeneous cellular identities emerge via a population-based inhomogeneous steady state solution

We consider a generic case where the single cell dynamics is governed by a minimal model of a genetic toggle switch, composed of two genes *u* and *v* that inhibit each other’s transcription via their respective promoters P_u_ and P_v_. We assume here that the extracellular signaling molecules *s* negatively affect the transcription of the switch gene *u* via intracellular signaling processes, while their expression is in turn regulated by the dynamics of the toggle-switch (Eq. (1), Fig. 1a, inset). As the communicating signals are secreted by the cells themselves, their concentration is no longer a parameter, but rather a variable in the system. This couples the states of the cells, creating interdependence between them, thus effectively establishing a single joint dynamical system.

**Fig. 1.**
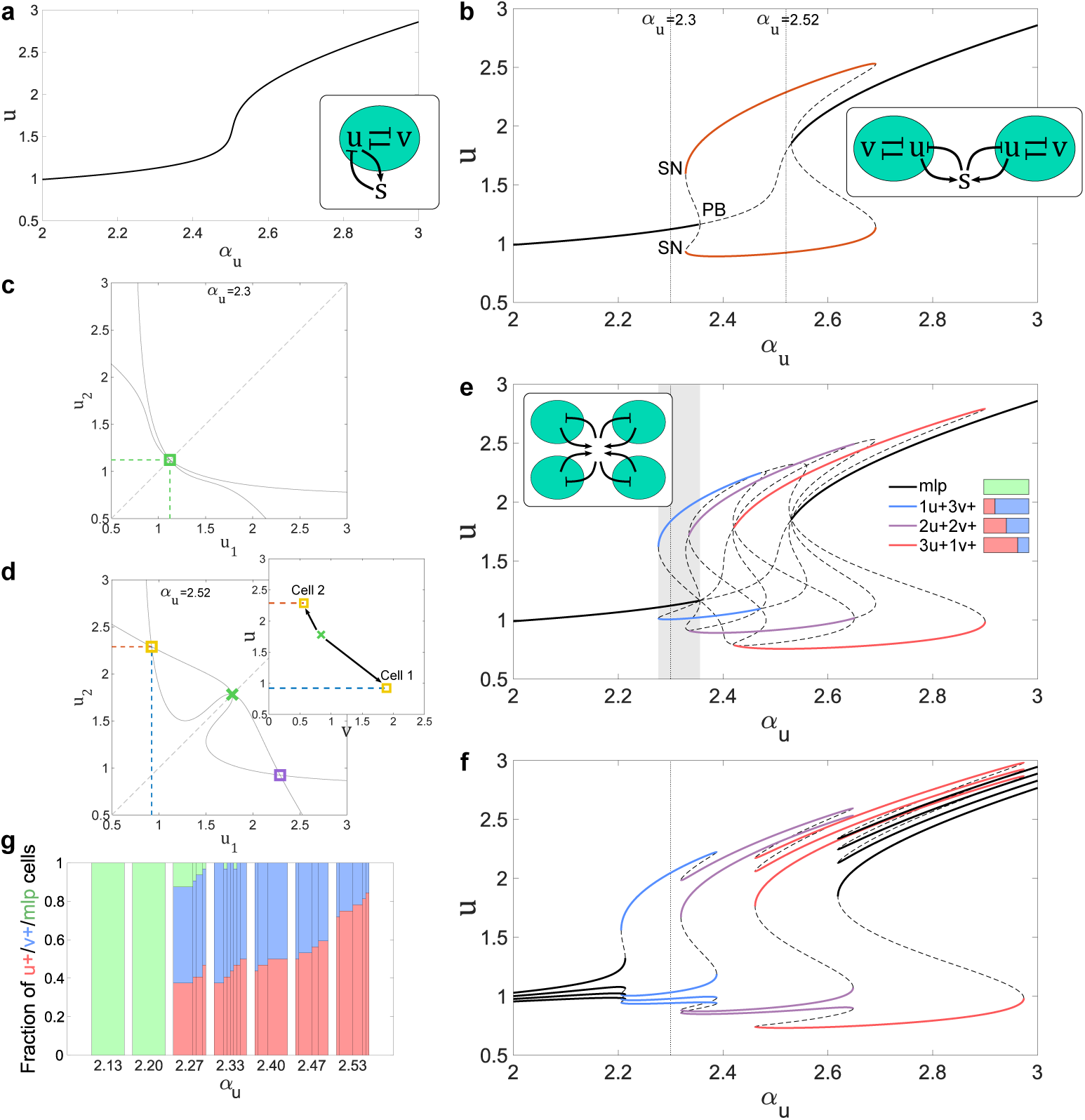
Emergence of cellular identities via population-based symmetry breaking. **a** Bifurcation diagram depicting monostable increase in *u* levels with respect to changes of promoter strength *α*_*u*_, in a single-cell system. Inset: underlying network topology. **b** Bifurcation analysis for two coupled identical cells (scheme in inset) reveals emergence of inhomogeneous steady state (IHSS) solution. PB: symmetry-breaking subcritical pitchfork bifurcation; SN: saddle-node bifurcation. Solid lines depict stable: homogeneous steady state (HSS, black) and IHSS (red); Dashed lines: unstable steady states; Dotted lines: organization points in parameter space. **c** *u*_1_-*u*_2_ phase plane analysis for organization of the two-cell system in the HSS (*α*_*u*_ = 2.3); solid lines: nullclines. **d** *u*_1_-*u*_2_ phase plane analysis for organization of the two-cell system in the IHSS (*α*_*u*_ = 2.52). Inset: *u*-*v* phase plane manifestation of the IHSS on the marginal level of single cells. **e** Bifurcation analysis of a system of *N* = 4 globally-coupled cells (inset). 3 stable IHSS distributions with increasing *u*+/*v*+ cell ratios appear sequentially (legend). Gray shaded area: HSS/IHSS coexistence parameter range for subcritical organization. Line description as in **b**. **f** Bifurcation analysis for *N* = 4 non-identical globally-coupled cells (equivalent as in **e**, see Methods). **g** *u*+/*v*+/mlp cell proportions for increasing *α*_*u*_, when a non-locally coupled population of *N* = 32 cells on a 4 *×* 8 grid is considered. The width of each sub-bar within a bar reflects the fraction of occurrence of the respective *u*+/*v*+/mlp proportion in the 10 independent realizations. The initial conditions were randomly drawn from a normal distribution 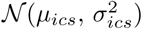 around the corresponding *α*_*u*_-specific mlp state as mean (*µ*_*ics*_), and *σ*_*ics*_=0.1. Model parameters in Methods.

The dynamics of the system on the level of a single cell, explored with respect to changes in the expression strength of the promoter P_u_, *α*_*u*_, exhibits monostability in the full parameter range (Fig. 1a). However, the bifurcation analysis for coupled systems revealed multiple different dynamical regimes, even for a minimal population of two identical cells. For a given range of *α*_*u*_, only a single fixed point is stable (Fig. 1b and Fig. 1c). This homogeneous steady state (HSS) represents the multilineage primed state (mlp), where both *u* and *v* are co-expressed in all cells. At a critical *α*_*u*_ value, the HSS’s symmetry is broken via a pitchfork bifurcation (PB): the HSS looses its stability, and a pair of fixed points is stabilized, giving rise to an inhomogeneous steady state (IHSS) (Fig. 1b, red). The IHSS is a single dynamical solution that has a heterogeneous manifestation: the unstable HSS splits into two symmetric branches that gain stability via saddle-node (SN) bifurcations (Koseska et al., 2013), and correspond to a high *u*-expression state in one cell (*u*_2_) and low *u*-expression state in the other cell (*u*_1_ *< u*_2_, yellow) and vice versa (violet, Fig. 1d). These branches reflect the differentiated cellular identities that emerge from the multilineage primed HSS. The results therefore show that tristability in single cells, i.e. initial coexistence of all the possible attractors, is not a necessary requirement to describe differentiation from a multilineage primed state, but rather emerging cellular fates are generated with coupling cell-cell interactions.The observations are furthermore valid for number of different network topologies (Supplementary Figs. 1a to 1c, Eqs. (2)).

Unlike the independent steady states in a classical bistable system, in the two IHSS branches the cell states are interdependent, and the branches are conjugate to one another. For two coupled cells therefore, a high *u*-expressing state in one and a low *u*-expressing state in the other cell always emerge jointly, even when the cells are completely identical based on their parameters. In that sense, the IHSS arising via a pitchfork bifurcation is a true symmetry breaking solution, since it inevitably and robustly leads to differentiated cellular identities. More generally, for *N* globally coupled cells, *N -* 1 different distributions of the cells are possible between the upper and the lower branches, manifested by stable attractors in phase space (Koseska et al., 2010). For example, for *N* = 4 globally coupled cells, three different IHSS distributions are stable: 1*u*+3*v*+ denotes one cell having high-(*u*+) and 3 cells having low-expressing *u* state (v+), 2*u*+2*v*+ denotes 2 cells in each state and 3*u*+1*v*+ denotes 3 cells with high- and one with low-expressing *u* state (Fig. 1e). These distributions span the parameter space and are always sequentially ordered towards increasing number of cells with high-*u* expression state for increasing *α*_*u*_, where branches of neighbouring distributions with similar proportions overlap in parameter space. Thus, it follows that reliable proportions of cells in the differentiated fates is an inherent property of the IHSS solution. Which proportion will be observed for a specific system only relies on the organization of that system in parameter space.

Parameter differences between single cells on the other hand increase the stability region of the IHSS (Koseska et al., 2009). Bifurcation analysis in the case where cell-to-cell variability in *α*_*u*_ was present (Methods) revealed that the HSS is already characterized with a slightly different mlp value in each cell, whereas its stability interval was relatively decreased (black, Fig. 1f). The parameter range where the IHSS solution is stable was on the other hand further expanded, and the overlapping intervals between neighbouring distribution branches were reduced (compare relative decrease from Fig. 1e to Fig. 1f). This effectively increases the robustness of the cell proportions in the two differentiated fates for a given *α*_*u*_. Thus, the number of cells acquiring specific fates is conserved under cell-to-cell variability.

These results demonstrate that several crucial dynamical characteristics emerge for a population-based symmetry-breaking: i) description of the undifferentiated co-expression state does not require evoking higher-order multistability on the level of a single cell, ii) heterogeneous cellular identities are generated from initially identical cells and are maintained on a population level due to cell-cell communication and iii) reliable proportions of differentiated cells are a direct consequence of the IHSS solution and can be robustly maintained under cell-to-cell variability.

To probe whether these principles generally apply for signaling with different communication ranges, we considered in addition to the global (all-to-all) coupling, also three different cell-cell coupling scenarios: local (nearest neighbor only), non-local (nearest and second nearest neighbour) communication, as well as distance-dependent cell-cell interactions forming an irregular grid (Supplementary Figs. 1d to 1g). The lattest was implemented by a coupling scheme where the probability of interaction declines with increasing distance between the cells (Supplementary Fig. 1f, left, Methods). Although the HSS was destabilized for different *α*_*u*_ values depending on the coupling type, the proportion of high-*u* expressing cells progressively increased with increasing *α*_*u*_ for non-locally, globally and irregularly coupled *N* = 32 cells on 4 *×* 8 grid (Fig. 1g, Supplementary Figs. 1d and 1f), as predicted by the progression of branches in the generic bifurcation analysis for globally coupled cells (*N* = 4, Fig. 1e). Multiple realizations starting from distributed initial conditions (Methods) demonstrated that although neighbouring IHSS distributions were populated, *α*_*u*_-specific cell type ratios still remained reliable with a deviation of ≲ 10%. For local coupling however, 50-50% ratio was maintained in a large *α*_*u*_ interval, indicating that visiting an IHSS manifestation that is different from a regular salt-and-pepper pattern on a 4*×*8 lattice only increases for higher *α*_*u*_ values (Supplementary Fig. 1e). This analysis however also highlighted that for *N* = 32 cells, specifications were observed for *α*_*u*_ values for which in the case of *N* = 2 coupled cells, only the multilineage primed state (the HSS) was stable (compare Fig. 1g and Fig. 1b). This inevitably opens the question how the timing of cellular differentiation comes about.

### Timing of cellular differentiation emerges in self-organized manner for critically organized growing populations

It is typically assumed that a change of a bifurcation parameter, such as extracellular concentration of signaling molecules *s* for example, drives the system through a dynamical transition, thereby relating the onset of differentiation to characteristic reaction rates (De Mot et al., 2016). Considering on one hand that *s* is not a parameter but a dynamical variable of the system, and on the other hand that the system parameters such as the promoter expression strength in the studied example are biochemical constants that cannot significantly deviate, question has to be posed what triggers the symmetry breaking event during differentiation. We therefore study next how cell fate specification from the multilineage primed state occurs for fixed system organization in parameter space.

Experimental observations show that the multilineage primed state is maintained for several cell cycles before differentiation occurs (Saiz et al., 2016; Hatakeyama et al., 2004). Given this initial symmetry, it follows that for *N* = 2 coupled cells the system is poised in the HSS, before the pitchfork bifurcation (as for *α*_*u*_ = 2.3, Fig. 1b). Since for *N* globally coupled cells, *N -* 1 IHSS distinct distributions are possible (as in Fig. 1e), it can be deduced that the number of distributions increases as 2^*n*^, where *n* denotes the step in the lineage tree. As the IHSS emerges via a subcritical PB and thereby coexists with the HSS in the vicinity of the PB point, inclusion of these new distributions with each cell division widens the coexistence regions in parameter space, eventually capturing the system’s organization point (compare gray shaded area in Fig. 1e to Fig. 1b). As a result, in the parameter region where for *N* = 2 coupled cells only the HSS was stable (Fig. 2a, green), for *N* = 4 non-locally coupled cells, stable IHSS solutions appeared (red/blue *u*+/*v*+ stackbar markers, Methods). Furthermore, for growing populations of *N >* 4 cells, the HSS loses its stability as the PB position is shifted due to an increase in the intrinsic spatial inhomogeneities, arising from the differing number of neighbors near the boundaries of the non-locally coupled grid. This shift is alike the one observed previously when cell-to-cell variability was introduced in the globally coupled system, reducing the HSS stability range (Fig. 1f). Therefore, while the PB position in a globally-coupled system of identical cells does not change (dotted line), for smaller communication range reflecting non-locally-(solid line) and locally-coupled (dashed line) systems, the symmetry breaking point shifts towards lower *α*_*u*_ value with cell number increase (Fig. 2b). This two parameter bifurcation diagram therefore shows that differentiation timing is not constrained to a narrow parameter region, thereby reducing the necessity for fine-tuning in locally and non-locally coupled systems. Taken together, the coexistence between IHSS and HSS and subsequent loss of HSS stability with increase in system size effectively generates a subcritical pitchfork bifurcation with respect to *N* (Fig. 2a). This renders the number of cells an effective bifurcation parameter that drives the triggering of symmetry breaking and cellular differentiation.

**Fig. 2.**
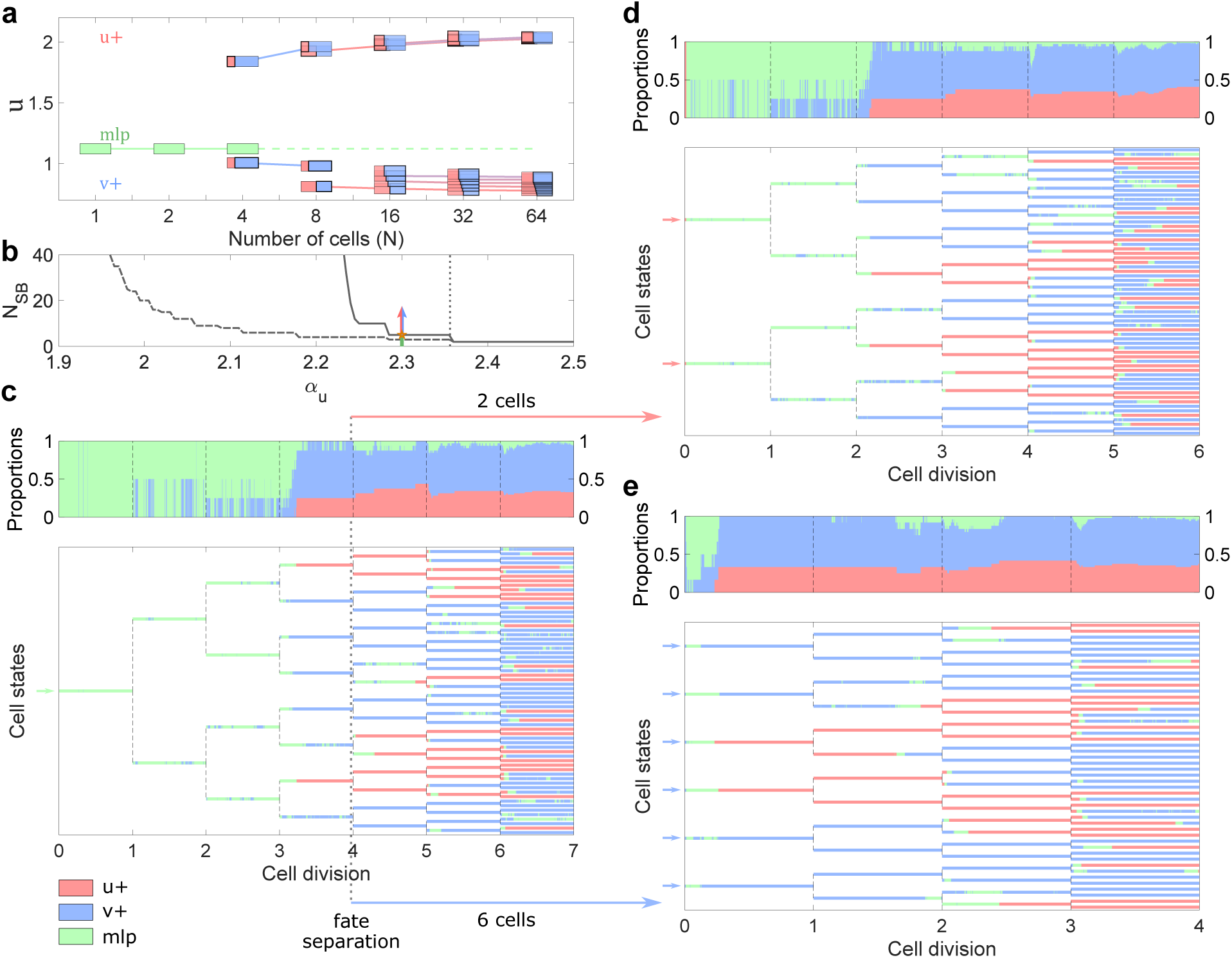
Critical organization before the pitchfork bifurcation point determines timing of differentiation events. **a** Emergence of stable IHSS distributions for increasing cell number *N* and organization before the pitchfork bifurcation point (*α*_*u*_ = 2.3). Existence of attractors with distinct *u*+/*v*+/mlp distributions is estimated with exhaustive stochastic search of the non-locally coupled system (see Methods). In analogy to Fig. 1e, depicted are the averaged *u*-values of the *u*+ cells (upper branch), and the corresponding *v*+ cells (lower branch) from each identified distinct distribution. Lines depict continuations of distributions with same proportions. Green: HSS mlp; red/blue stackbar markers: *u*+/*v*+ proportions in the IHSS solutions. Other parameters as in Fig. 1b. **b** Population size threshold at which symmetry is broken (*N*_*SB*_) in relation to *α*_*u*_. Dashed line: *N*_*SB*_ for local coupling; solid line: *N*_*SB*_ for non-local coupling; dotted line: fixed position of PB for global coupling. **c** Lineage tree exhibiting characteristic differentiation from the multilineage primed- to the *u*+/*v*+ states as a function of the cell number. System’s organization as in **a**. Green: cells in the multilineage primed state; Red/blue: cells with high/low *u*-expression state (*u*+/*v*+). Upper panel: respective cell type proportions. **d**, **e** Lineage trees generated from homogeneous sub-populations of differentiated cells. At *n* = 4th step of the lineage tree (*N* = 8 cells) in **c**, differentiated cells of identical fates are separated, and the respective evolution of the sub-systems is followed to generate: **d** Lineage tree seeded from the *N* = 2 cells that before separation adopted the high *u*-expression state (*u*+); **e** Lineage tree seeded from the *N* = 6 cells that initially had adopted the low *u*-expression state (*v*+). Both upper panels reflect the respective cell type proportions.

To demonstrate how organization before the bifurcation point in conjunction with cell division can serve as a timing mechanism that regulates the onset of differentiation, we generated a lineage tree where the population growth of non-locally coupled cells on a grid is represented in a simplified manner: after a given time period T that mimics cell cycle length, all cells divide and the number of cells is doubled. The initial gene expression states of the daughter cells are inherited from the final state of the mother cell. Starting from the mlp state (green), as the cell population grows in size and the IHSS distributions appear via the subcritical PB, the loss of HSS stability triggers switch to the already existing IHSS solution (*n* = 4th step of the lineage tree, Fig. 2c). The distribution proportions for increasing system size showed a steady ratio above a certain population size (*N* ≈ 16 cells, Fig. 2c, upper panel), in agreement with Fig. 1g.

The self-organized manner of generating heterogeneous cellular identities on a population level that we propose here also implies that differentiated fates cooperatively coexist. To check the importance of cooperativity to maintain cell fates, we performed a numerical experiment in which cells at *n* = 4th step of the lineage tree (*N* = 8 cells, Fig. 2c) are separated according to their fates, forming two single-fate sub-populations of different size that can further continue to grow and divide (Figs. 2d and 2d). The sub-population of two coupled cells with high *u*-expression reverted to the only stable 2-cell system attractor – the mlp HSS, but after 2 cell cycles (*N* = 8 cells) both differentiated fates re-emerged (Fig. 2d). The other sub-population of *N* = 6 cells with low *u*-expression on the other hand, initially briefly re-visited the multilineage primed state before both cell types stably re-emerged and the population settled in the IHSS attractor (Fig. 2e). The difference in timing between the two cases again points to the cell number dependence in triggering of the symmetry breaking (Figs. 2a and 2b). The cell type ratios for both sub-populations of different size were stabilized to a steady value similar to that of the full system before separation, and differed among each other by around 6%. This scaling and regenerating capability of the self-organizing system is a direct consequence of the properties of the IHSS solution: dynamically, it is not permitted to populate the upper without populating the lower *u*-expressing state. Thus, even when cells are separated such that only the cells whose dynamics with high (low) *u*-expression state remain, the cell division and the cell-cell communication through which IHSS is established in a first place, will enable the system to recover both cell types with reliable ratios.

### Reliable proportions of differentiated cells are an intrinsic feature of the IHSS solution

To systematically probe the robustness of cell type ratios, we investigated the effects of variation in the initial conditions, as well as presence of intrinsic noise that is ubiquitous in gene expression. Results are obtained for a population of *N* = 32 cells under the 4 distinct coupling types, and a fixed *α*_*u*_ organization before the symmetry breaking bifurcation as in Fig. 2c (*α*_*u*_ = 2.3). Sampling the single cell initial conditions from normal distributions with increasing standard deviations *σ*_*ics*_ around the mlp-value produced distributions with reliably conserved proportions between *u*+ and *v*+ cells for each coupling type. The different communication ranges yielded different stable *u*+/*v*+ proportions for this fixed *α*_*u*_ value, in agreement with values in Supplementary Fig. 1, Fig. 1: *∼*0.45 for non-local coupling (Fig. 3a), 0.5 salt-and-pepper patterns for local coupling (Supplementary Fig. 2a), *∼*0.4 for irregular coupling (Supplementary Fig. 2g), whereas the HSS remained stable against moderate perturbations for global coupling (Supplementary Fig. 2d). Stochastic realizations with gradual shift in the initial mean value from high *v*-expression to high *u*-expression state (*µ*_*ics*_ from 0 to 1, Fig. 3b, Supplementary Figs. 2b, 2e and 2h), or with increasing noise intensity (Fig. 3c, Supplementary Figs. 2c, 2f and 2i), also demonstrated reliable *u*+/*v*+ proportions. As observed previously, organization in parameter space for a given communication range determines the obtained value of steady proportions of cells in the differentiated fates. We observed as well a manifestation where besides populating *u*+/*v*+ states within the IHSS solution, few cells also populated the mlp state, resembling a chimera-like state (Kuramoto and Battogtokh, 2002; Abrams and Strogatz, 2004). This was however only observed for non-local or irregular coupling (Fig. 3a, Supplementary Figs. 2g to 2i). Since chimera states have been predominantly characterized for systems of coupled oscillators, a detailed theoretical study is required to dynamically classify this solution.

**Fig. 3.**
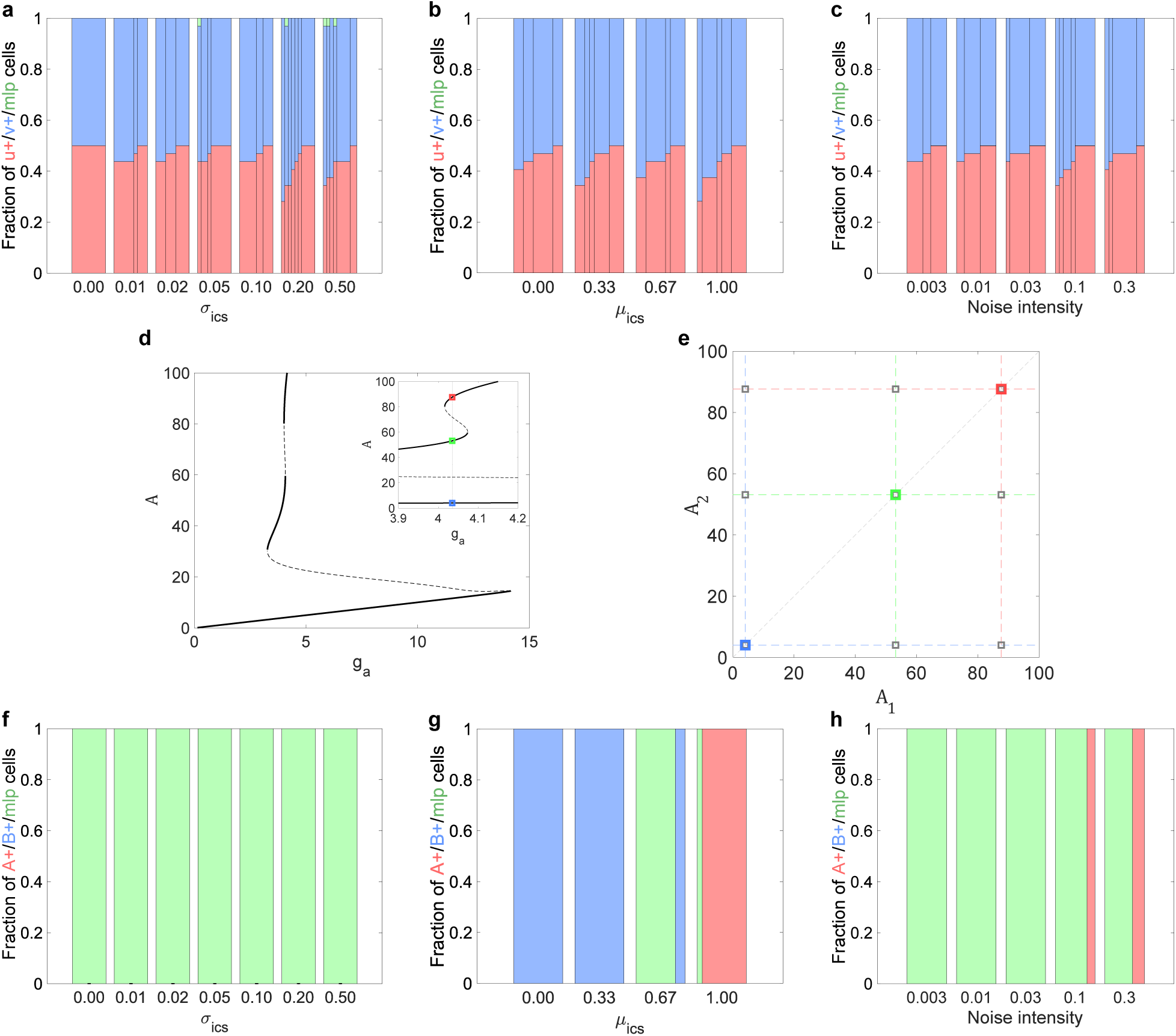
Comparison of robustness in cell ratios between symmetry-breaking and pre-existing asymmetry-based mechanisms. *u*+/*v*+/mlp cell type proportions for: **a** gradual increase in the standard deviation (*σ*_*ics*_) around the initial conditions’ mean mlp value; **b** gradual shift in the distribution’s mean (*µ*_*ics*_) initial value from high v-expression to high u-expression state for stochastic realizations; and **c** increase in the noise intensity. The bar graphs were calculated as in Fig. 1g, for *α*_*u*_ = 2.3 and *N* = 32 non-locally coupled cells. **d** Bifurcation diagram of the system in Eqs. (3) exhibiting tristability on a single-, as well as population-level (inset: magnified tristability region); **e** Attractor occupancy for both cases. Colored squares represent the three possible solutions of the coupled 2-cell system, whereas gray squares depict the possible combinations of single cell solutions. **f, g, h** *u*+/*v*+/mlp cell type proportions (conditions as in **a, b, c**, respectively) for *N* = 32 non-locally coupled cells exhibiting tristability on a population level (Eq. (3)).

We analyzed next whether reliable cell proportions can be achieved when considering that cellular identity is an intrinsic feature of single cells. Following this current view of differentiation, tristability on the level of single cells is the necessary requirement to simultaneously account for the multilineage primed as well as the differentiated fates, where intercellular signaling only drives the distribution of cells in one of the existing attractors (De Mot et al., 2016). It has been shown however that cell-cell communication can lead to novel dynamical solutions of the coupled system that are different than those of the isolated cells, indicating that the features of the coupled system cannot be formally explained with those of single cells (Suzuki et al., 2011; Goto and Kaneko, 2013; Koseska et al., 2007; Ullner et al., 2008). The proposed symmetry breaking mechanism is also a demonstration of this principle. We therefore explore whether the concept of multistability, on the level of a single-cell as well as on a joint communicating system level, is necessary and sufficient to serve as a basis for differentiation.

For this purpose, we adapted a paradigmatic multistability model where toggle switch with self-activation leads to tristability on the level of single cells (Jia et al., 2017), to also exhibit tristable behavior when cells are coupled (Eq. (3), Fig. 3d). In this case, the linage commitment was not robust, and the fate decisions were completely determined by the distributions of initial conditions or the amplitude of the noise intensity. All cells either remained in the multilineage primed state, or all cells jointly populated one of the two differentiated states (Fig. 3e, Figs. 3f to 3h). Thus, while a system of independent cells can theoretically populate any combination of the coexisting single-cell attractors (Fig. 3e, gray squares), within a coupled system all of the cells populate the same state in the multistability solutions (Fig. 3e, coloured squares). This comes from the lack of symmetry breaking (PB) (Fig. 3d) which highlights the difference in dynamical behavior to the IHSS solution: the IHSS arises from a true symmetry breaking solution, its branches emerge together at the PB and as previously noted, are conjugate to each other (van Kekem and Sterk, 2019), always leading to a joint occupancy of each of the differentiated fates by the cells.

Existence of multistability on a single-cell level is therefore not a sufficient condition for co-existence of heterogeneous cellular identities and reliable proportions on a population level of a coupled system. While in principle coupled multistable systems could generate a more complex PB-induced IHSS solution, as we have described here they are not a necessary condition for symmetry breaking emergence.

### IHSS as a generic mechanism for cellular differentiation: examples of the early embryo and vertebrate neurogenesis

The proposed symmetry breaking solution together with the system’s organization before the pitchfork bifurcation point uniquely provides a dynamical mechanism of differentiation that simultaneously accounts for robustness in proportions and self-organized timing of the event. These properties can be directly related to the population-based dynamical transitions and therefore they apply to systems with diverse gene expression dynamics in single cells. In addition to the generic model that displays monostability in a single cell system, we demonstrate this using bistable and oscillatory single cell behavior, as pervasive during embryogenesis and neurogenesis, respectively.

During early embryogenesis, differentiation from the ICM state results in the formation of Nanog-positive epiblast (Epi) and Gata6-positive primitive endoderm (PrE) cells. Multistability on a single cell level has been proposed as an underlying mechanism, where cell fate specification is mediated by intercellular interactions involving Fgf4 communication and Erk signaling (Schröter et al., 2015; Bessonnard et al., 2014; Chazaud et al., 2006). The heterogeneities in extracellular Fgf4 concentrations that each cell perceives have been determined as crucial for cells to populate one of the remaining stable attractors (De Mot et al., 2016; Bessonnard et al., 2014). This single-cell identity view has also been used to explain the occurrence of purely Epi or PrE states when development occurs in the absence of Fgf4 signaling or in presence of a constant high level of exogenous Fgf4 (Nichols et al., 2009; Yamanaka et al., 2010).

Considering again that Fgf4 is not a parameter, but rather a variable in the system, we studied the dynamical properties of a minimal model of two coupled cells (Eq. (4)), where the single-cell dynamics is characterized with a bistable behavior (Schröter et al., 2015). Bifurcation analysis demonstrated that not only a co-expression ICM-like state (black) emerges due to cell-cell communication, but also a symmetry breaking IHSS occurs, reflecting the differentiation in PrE and Epi fates (red, Fig. 4a). Numerical simulations of a population of *N* = 32 locally-coupled cells organized before the pitchfork bifurcation (*α*_*N*_ = 5) showed the transition of the joint system towards either Epi or PrE fates upon signaling perturbations, as experimentally observed (Nichols et al., 2009; Yamanaka et al., 2010). Administering increasing doses of Fgf4 inhibitor gradually decreased the cell-cell communication, eventually unravelling the Nanog+ solution of the single-cell system (Fig. 4a, gray profile). Thus, gradual increase of the Epi/PrE proportions towards an all-Epi state was observed (Fig. 4b). On the other hand, increasing the dose of exogenous Fgf4 overwrote the intercellular communication such that the system reflected the cells’ dose-response behavior, resulting in an abrupt joint switch in the population to high Gata6 expression. The coupled system either remained in the salt-and-pepper pattern or transitioned jointly to the PrE state (Fig. 4c), in line with the experimental observations in (Kang et al., 2013). Taken together, these results suggest that IHSS in conjunction with critical organization before the pitchfork bifurcation is consistent with the existing experimental observations and thus, this general principle of population-based symmetry breaking can serve as a basis for robust Epi and PrE specification, crucial during early development.

**Fig. 4.**
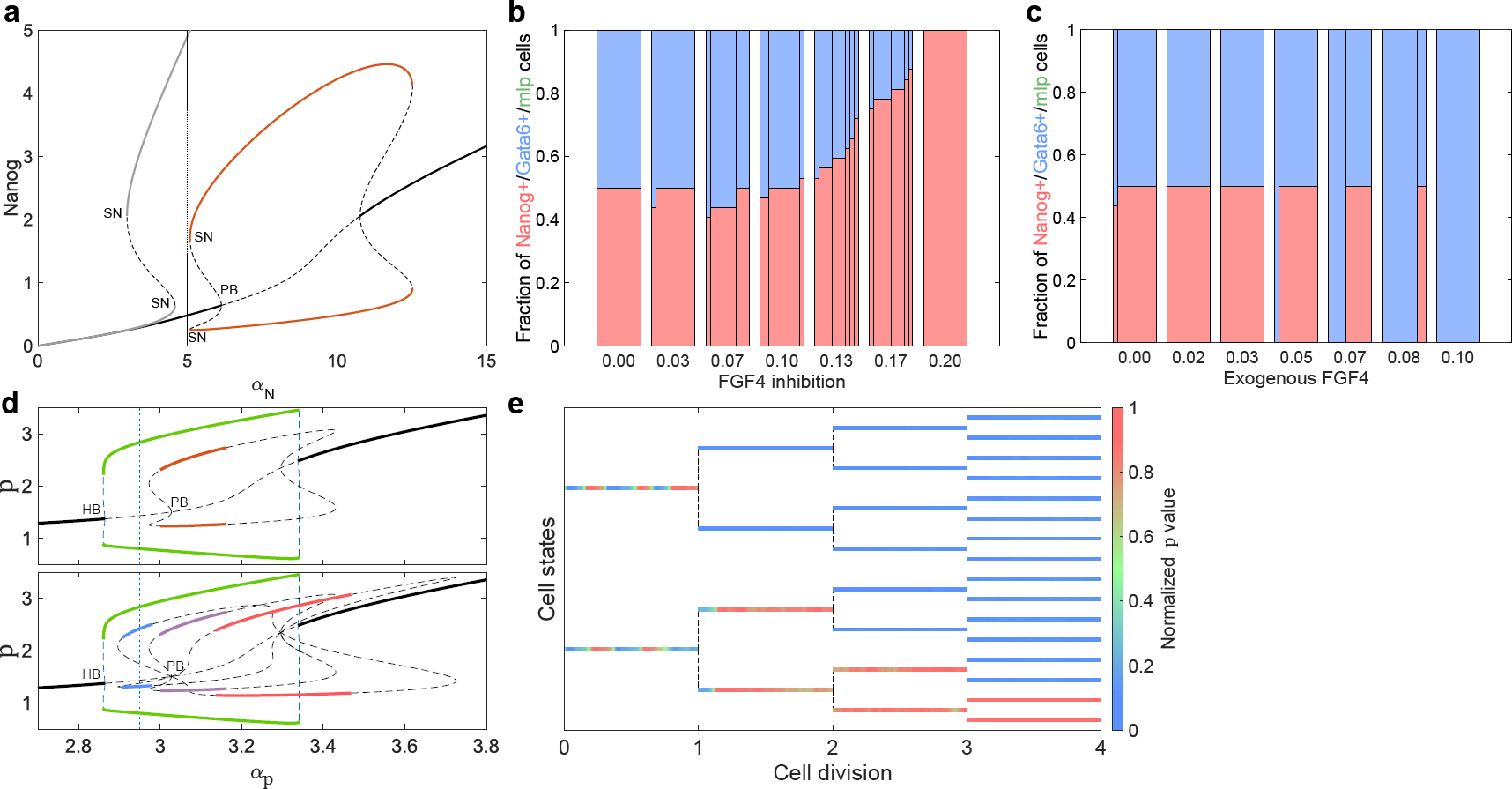
Symmetry-breaking via pitchfork bifurcation as a generic mechanism for differentiation during early embryogenesis and neurogenesis. **a** Combined bifurcation plot depicting bistable Nanog-Gata6 behavior on the single-cell level (gray profile) and IHSS on a 2-cell population level (red), (Eq. (4)). Line description as in Fig. 1b. **b, c** Administering increasing doses of Fgf4 inhibitor (**b**) or exogenous Fgf4 (**c**) leads towards increasing occupancy of Epi and PrE fates, respectively. **d** Bifurcation diagram for *N* = 2 (top) and *N* = 4 (bottom) globally coupled paradigmatic oscillators, mimicking early neurogenesis. Solid lines denote stable attractors: HSS (black), homogeneous stable limit cycle denoting synchronized oscillations (green), IHSS (red, top), and different IHSS distributions (bottom): blue – 1*p*+3*q*+ (1 high *p*-state, 3 low *p*-state cells), violet – 2*p*+2*q*+, red – 3*p*+1*q*+. Dashed lines depict unstable: steady states (black), limit cycle (blue). **e** Lineage tree demonstrating differentiation from a homogeneous cell population that displays synchronized oscillations (Eq. (5)) into two distinct differentiated fates as a function of the cell number. Stochastic simulations were performed as described in Methods.

In the developing mouse brain on the other hand, oscillatory expression of transmembrane ligands of the Delta, Serrate, and Lag-2 (DSL) family, Hes-Her proteins, and proneural proteins have been observed in neural precursors, before patterned steady states are reached (Kageyama et al., 2007; Shimojo et al., 2008; Momiji and Monk, 2009). Since the oscillations possibly play a central role in delaying the onset of neural differentiation, multiple models have already provided dynamical descriptions of these observations (Momiji and Monk, 2009). However, a dynamical mechanism that leads to heterogeneous steady state levels from an initially homogeneous oscillatory dynamics is lacking. Without describing the molecular details of the system, but rather using a paradigmatic model that captures oscillations on the level of single cells (Eq. (5), Methods), we next explore the possibility that the symmetry of the synchronized solution of the coupled system can be broken via a pitchfork bifurcation induced by the intercellular communication. The bifurcation analysis of a minimal (*N* = 2) globally coupled system showed a coexistence between a limit cycle solution corresponding to synchronized oscillations and an IHSS corresponding to the differentiated state (Fig. 4d, top). Increase in cell number, similarly as before (Fig. 1b to Fig. 1e), enlarged the parameter region where the IHSS is stable, and therefore in the parameter range where for *N* = 2 coupled cells only stable limit cycle was observed, for *N* = 4 cells stable IHSS distributions appeared. Numerical simulations where cell division was explicitly considered demonstrated that a transition from synchronized oscillations to a symmetry-broken IHSS emerges for critical organization before the pitchfork bifurcation (Fig. 4e). Even though the model does not reflect the molecular details of the Notch-pathway oscillatory dynamics during vertebrate neurogenesis, it demonstrates that population-based symmetry-breaking can in principle serve as a mechanism to describe differentiation for growing populations, characterized with oscillatory dynamics. Thus, using these two models that resemble early embryogenesis and neurogenesis cases, together with the generic examples (Fig. 1, Supplementary Figs. 1a to 1c), we demonstrate the general applicability of the proposed population-based symmetry breaking mechanism via a pitchfork bifurcation. Subcritical organization before the pitchfork bifurcation on the other hand again enables self-organized timing of cellular differentiation to emerge due to cellular division.

## Discussion

Important insights regarding symmetry breaking mechanisms unquestionably came from Turing’s seminal work (Turing, 1952), and have been subsequently widely explored to generally describe the emergence of spatial organization during development (Raspopovic et al., 2014; Economou et al., 2012). The population-based symmetry breaking mechanism we propose here however, not only provides a unique dynamical transition from homogeneous to a heterogeneous distribution of cellular fates, but in conjunction with subcritical organization enables accounting for the reliability and timing of the differentiation event. Although a similar mechanism of a population-based symmetry breaking via a pitchfork bifurcation has been previously suggested for the Delta-Notch lateral inhibition model when the strength of the interaction between the two cells is varied (Ferrell, 2012), it relied on a supercritical PB. Moreover, it has not been proposed how robustness and accuracy emerge due to this dynamical transition which to our understanding is more limiting since sub-critical organization, as we have demonstrated here, is crucial to determine timing of cellular differentiation in reliable cell type proportions.

We argue that intercellular communication, an integral part of developing mammalian embryos, gives rise to differentiated fates whose dynamical manifestation cannot be directly inferred from the gene regulatory network dynamics in single cells. We demonstrated that such novel dynamical solution with a heterogeneous manifestation within the population can emerge with cell division even from systems of single cells, characterized with monostability. This is distinct from systems that rely on pre-existing asymmetries where intercellular signaling is influencing the distribution of cellular fates between co-existing attractors.

In order to understand the basic mechanisms of cell-to-cell cooperative behavior, several theoretical principles have been already successfully developed by investigating both natural and synthetic genetic networks (McMillen et al., 2002; Taga and Bassler, 2003; Kuznetsov et al., 2004; Garcia-Ojalvo et al., 2004; Ullner et al., 2007). These studies have shown that multiple coexisting attractors arise in multicellular systems, thereby enabling a very diverse dynamics, different than the dynamics of the isolated cells, to be manifested on a population level in conjunction with high adaptability. Cooperative behavior has been therefore profiled as necessary for the emergence of typical features observed in cellular systems. In that respect, it has been proposed that phase-repulsive coupling generally leads to inhomogeneous solutions such as the IHSS discussed here, whereas the size of the system affects the relative sizes of the basins of attraction of the coexisting regimes to promote the inhomogeneous solutions (Ullner et al., 2008; Koseska et al., 2010). Another inhomogeneous solution that is directly associated to the IHSS is an inhomogeneous limit cycle – a periodic solution of a system of coupled oscillators rotating around two spatially non-uniform centers (Ullner et al., 2007, 2008; Koseska et al., 2010). Manifestation of this solution has been in turn related to stem cell differentiation with self-renewal (Suzuki et al., 2011). The derived generalizations how these solutions emerge however mainly refer to populations of coupled genetic networks that exhibit oscillations. In order to understand the minimal coupling principles that lead to novel inhomogeneous solutions in populations that exhibit mono- or bistability on a single cell level, a more detailed theoretical analysis is required. The hypothesis that timing is triggered by the number of cells can be experimentally tested for example in stem cell culture where the specification of populations of different sizes is tracked over time. Such experiments can also shed light on the self-organizing properties of mammalian embryos, by probing the regeneration of cell type proportions upon external physical or chemical perturbations, as suggested in Fig. 2. The co-operativity necessary for these organization principles to arise can be further tested in systems of coupled synthetic genetic networks that can mimic intercellular communication.

It is important to note here that one of the main characteristics of the IHSS solution is the robustness of the differentiated cell types numbers. The defined number of possible distributions between the two cell fates – all of which represent stable attractors, therefore ensures that stochasticity in gene expression dynamics or variability in the initial conditions can only switch the system between neighbouring attractors with similar proportions, thus preserving the overall robustness. This in turn allows to envision further extension of this principle of population-based symmetry breaking to describe pluri-/multipotency of stem cells. Conceptually, this would correspond to a finite cascade of subsequent pitchfork bifurcations occurring simultaneously on both branches of the existing IHSS solutions (Zakharova et al., 2013; van Kekem and Sterk, 2019).

Our results overall suggest that the cooperative behavior of growing populations enables the symmetry of a homogeneous population to be broken, as a pre-requisite for novel information regarding different cellular types to emerge, whereas organization in the vicinity of this dynamical transition allows to comprehensively capture how robustness and accuracy during development are generated.

## Methods

### Generic cell-cell communication system

The generic model from Figs. 1a to 3c and Supplementary Figs. 1d to 2i is described with the following set of equations:

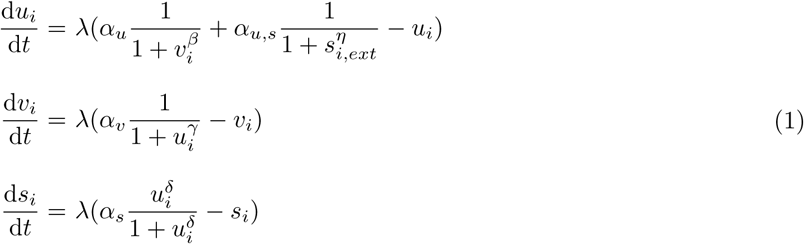

Here *u* and *v* are the two genetic markers that are coupled with mutual inhibition, while *s* is the secreted signaling molecule whose production is regulated by *u. i* – single cell index. In a single cell case, *s*_*i,ext*_ = *s*_*i*_, as in Fig. 1a. For multiple cells, the system is distributed spatially on a regular two-dimensional lattice with no-flux boundary conditions. Four different communication ranges, *R*, of the secreted signaling molecule were considered: globally connected network (all-to-all communication, *R* = *∞*), locally connected network (cells communicate only with direct neighbors on the lattice, *R* = 1*a*, where *a* is the lattice constant), non-locally connected network (cells communicate with direct neighbors and cells on two hops away on the lattice, *R* = 2*a*) and distance-based coupling on an irregular grid (cells communicate with other cells with probability 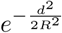, where *d* is the cell-cell distance and *R* = 1 in this case). In these cases 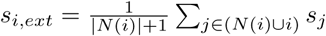 is the external amount of signal perceived by cell *i* from its neighborhood *N* (*i*), which negatively regulates the production of *u*. This effectively creates a joint 3*N*-dimensional system, where *N* is the total number of cells.

*α*_*u/v/s/u,s*_ are the production rates, while degradation rates are omitted as the system is globally scaled by *λ. β,γ,δ,η* are the Hill coefficients. Values of *α*_*u*_ = 2.3, *α*_*v*_ = 3.5, *α*_*s*_ = 2, *α*_*u,s*_ = 1, *β* = *γ* = *δ* = *η* = 2, *λ* = 50 were used throughout the study.

For the case of non-identical cells (Fig. 1f), the *α*_*u*_ parameter was uniformly varied between the cells in the range from *−*2% to 2% of its value.

For the stochastic simulations (Fig. 2, Fig. 3 and Supplementary Fig. 2), a stochastic differential equation model was constructed from Eq. (1) by adding a multiplicative noise term *σX*d*W*_*t*_, where d*W*_*t*_ is the Brownian motion term and *X* is the variable state. When the noise intensity *σ* was not varied, it was set to 0.1 was used unless when noise intensity was varied. The model was solved with Δ*t* = 0.01 using the Milstein method (Mil’shtein, 1974).

To discriminate between the multilineage-primed-(mlp), *u*-positive (*u*+) or *v*-positive (*v*+) cell fates for a given realization, each marginal cell state vector (*u*_*i*_,*v*_*i*_) within the converged state of the system (IHSS or HSS) was individually categorized as one of three fates and the three-term ratio (proportions) of the realization was subsequently calculated. The reference mlp fate vector was pre-determined for a given parameter set, i.g. for a specific value of *α*_*u*_, from the steady state value of 1-cell monostable system realization, since the bifurcation analysis demonstrated that the mlp HSS for a single cell and coupled systems are equivalent. Marginal cell state of a deterministic realization was categorized as mlp fate when its value fell within 1% around the pre-determined value, whereas cell states with larger *v*-value were *v*+, and with smaller *v*-value – *u*+ fates. Transient states for the stochastic realizations in Figs. 2c and 2d were categorized as mlp if they fell within 5% around the deterministic mlp state. End states of all stochastic realizations were allowed to converge to their deterministic attractor state in noise-free fashion before categorizing (with 1% std around mlp fate). Finally, results from 10 repetitions were grouped according to matching proportions and each proportion was plotted as a stacked sub-bar within a bar plot whose width corresponds to the number of repetitions in the group, i.e. the fraction of occurrence of that proportion in the simulations.

### Cell-cell communication systems exhibiting population-based symmetry breaking

To demonstrate the generality of the population-based symmetry-breaking mechanism via a pitchfork bifurcation, we also tested several minimal network topologies (Supplementary Figs. 1a to 1c). Their dynamics is described using the following systems of equations:

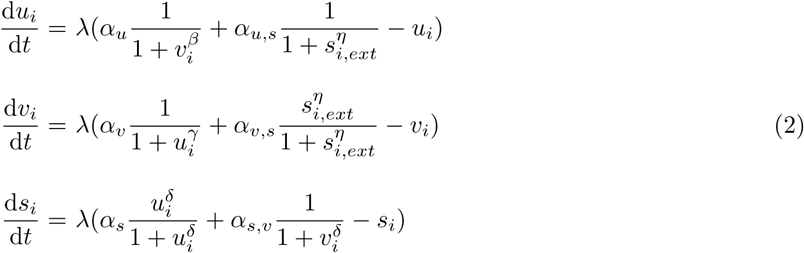

For Supplementary Fig. 1a: *α*_*u,s*_ = 0, *α*_*s*_ = 1, *α*_*v,s*_ = 1, *α*_*s,v*_ = 0 and *α*_*v*_ = 2.75; for Supplementary Fig. 1b: *α*_*u,s*_ = 0.5, *α*_*s*_ = 0, *α*_*v,s*_ = 0, *α*_*s,v*_ = 3 and *α*_*v*_ = 3; and for Supplementary Fig. 1c: *α*_*u,s*_ = 0, *α*_*s*_ = 0, *α*_*v,s*_ = 1, *α*_*s,v*_ = 2 and *α*_*v*_ = 3.

### Estimating IHSS distributions as a function of the number of cells (N)

By analogy to Fig. 1e, the different branches of the IHSS (i.e. proportions of cells in them) were estimated using the number of cells as a bifurcation parameter. For this, exhaustive scanning was performed to locate the different fixed point attractors in phase space for each *N*. The scanning process involved 20 repetitive executions with different noise intensities (varying from 0 to 0.3). Each repetition consisted of 30 alternating cycles of stochastic-(for exploration), followed by deterministic execution (for convergence to attractor), when the reached state was tested for stability and subsequently recorded. For every distinctly detected attractor, the *u* + */v* + */mlp* proportion of cells was estimated, after which the average *u*-value was calculated and plotted for each of the branches (the *u*+ and *v*+ cells for IHSS, or mlp cells for HSS; color-coded, see Fig. 2a). Chimera-like states were omitted from the diagram for brevity.

### Lineage tree generation

Generation of lineage trees was performed using stochastic simulations where the system doubles in size at regular time intervals, starting from a single cell system to an 8 *×* 8-cell system. At every cell division the mother cell’s steady state is passes on to daughter cells’ initial conditions. Cell divisions occur along horizontal and vertical axis on the grid alternately, sequentially yielding lattices of 1 *×* 1, 1 *×* 2, 2 *×* 2, 2 *×* 4, 4 *×* 4, 4 *×* 8 and 8 *×* 8. Cellular states were categorized in every time instance to plot the single cell temporal evolutions in the lineage trees (Figs. 2c to 2e). Further, cellular proportions in the system were estimated from those values, and their temporal evolution was shown in the panels above the lineage trees.

In the cell fate separation case (Figs. 2d and 2e), the steady states of the cells at the end of the fourth cycle were categorized and the differentiated cells were then separated: *u*+ cells were given as seeds to a new lineage tree (1 *×* 2 grid), while *v*+ cells were seeds for a separate one (2 *×* 3 grid). Following this, multiple cell divisions were again performed and the cell proportions were estimated.

### Multistability model on a single-cell/population level

Following (Jia et al., 2017) that demonstrated tristability on a single cell level, we introduced cell-cell communication to achieve tristability on a population level (Figs. 3d and 3e) with the following equations:

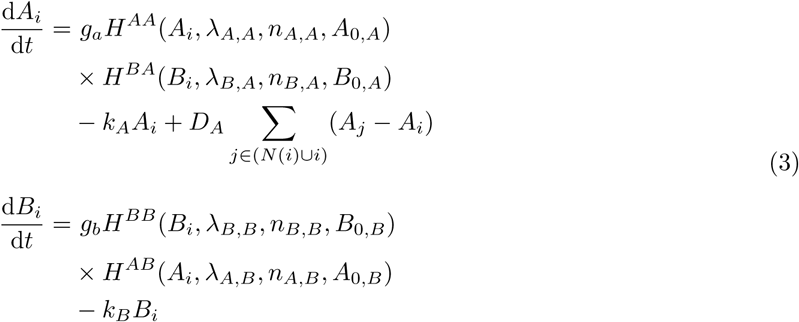

where the shifted Hill function is used to capture regulation of production of X by Y:

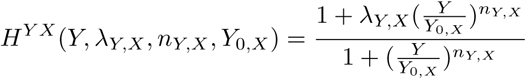

Cells communicate via diffusive coupling of *A* (*D*_*A*_ = 0.5). Other parameters: *λ*_*A,A*_ = *λ*_*B,B*_ = 3, *λ*_*A,B*_ = *λ*_*B,A*_ = 0.1, *A*_0,*A*_ = *B*_0,*B*_ = 80, *A*_0,*B*_ = *B*_0,*A*_ = 20, *n*_*A,A*_ = *n*_*B,A*_ = *n*_*B,B*_ = *n*_*A,B*_ = 4, *k*_*A*_ = *k*_*B*_ = 0.1, *g*_*b*_ = 5. *g*_*a*_ = 4.035 was set to place the system in tristability regime.

### Modeling of the Nanog-Gata6 differentiation system in the early embryo

Similarly to the generic *u*-*v* symmetry breaking system, the Nanog-Gata6 system was modeled using the following dynamics:

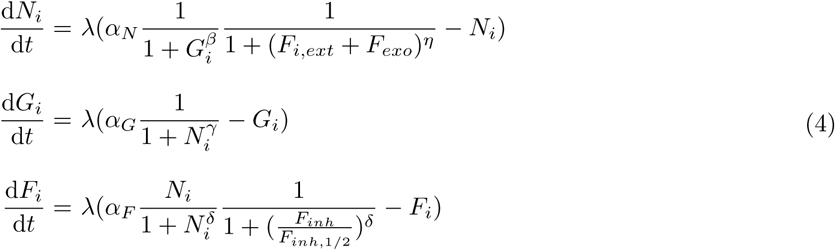

Here *N, G* and *F* are Nanog, Gata6 and Fgf4, respectively. *α*_*N*_ = 5 to place the system before the first bifurcation point (Fig. 4a) by analogy to the *u*-*v* system, 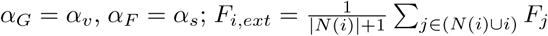; *F*_*inh*,1*/*2_ = 0.1 for the cases with external inhibition *F*_*inh*_ of Fgf4 and *F*_*exo*_ is the exogenous Fgf4 concentration. The other parameters were as in the *u*-*v* system. Locally connected network was used for the corresponding stochastic simulations with multiplicative noise term and *σ* = 0.5.

### Paradigmatic model mimicking the vertebrate neurogenesis process

It has been previously demonstrated that the presence of time delays in models of lateral inhibition can result in significant oscillatory transients before patterned steady states are reached. The impact of local feedback loops in a model of lateral inhibition based on the Notch signaling pathway, elucidating the roles of intra- and intercellular delays in controlling the overall system behavior have been also proposed (Momiji and Monk, 2009). Here, we aim at understanding whether population-based pitchfork bifurcation can provide the dynamical background behind the observed symmetry-breaking phenomenon. Since our aim is demonstrating the validity of this concept, we omit the molecular details of the Notch-pathway example and model a generic case where the gene expression dynamics in each cell is characterized with oscillatory behavior, whereas intercellular communication between the cells is realized in a global manner, for simplicity. The dynamics of the system is therefore described as:

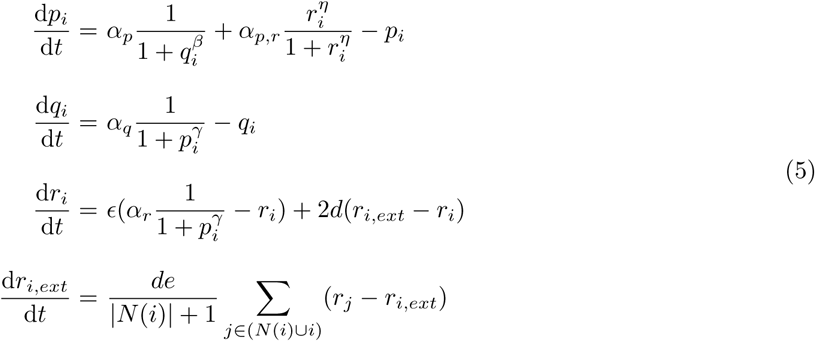

Here *p* and *q* are two genes that mutually inhibit each other’s expression, *r* controls the production of the signaling molecule, whose extracellular concentration is denoted as *r*_*ext*_. This system has been demonstrated to exhibit synchronized oscillations in a population of communicating cells (Kuznetsov et al., 2004; Koseska et al., 2007). *α*_*p*_ = 2.95 to place the system in the region where the limit cycle is stable (*N* = 2), and subsequently within the stability region of the IHSS for *N* = 4 (Fig. 4d). Other parameter values: *α*_*q*_ = 5, *α*_*p,r*_ = 1, *α*_*r*_ = 4, *β* = 2, *γ* = 2, *δ* = 2, *η* = 2, *ϵ* = 0.01, *d* = 0.008, *de* = 1. The lineage tree was generated by stochastic simulations (executed in C) with additive noise term of *σ* = 0.0008.

The numerical bifurcation analysis was performed using the XPP/AUTO software (Ermentrout, 2016).

All simulation except where explicitly noted were performed using custom-made code in MATLAB (MATLAB and Statistics Toolbox Release R2018b, The MathWorks, Inc., Natick, Massachusetts, United States).

## Acknowledgements

The authors thank Philippe Bastiaens for numerous discussions, as well as for critically reading the manuscript.

## Author contributions

A.K. conceptualized the study, A.S. performed the numerical simulations and bifurcation analysis with help of A.K., and C.S provided valuable insights on the molecular details of early differentiation. A.K. wrote the manuscript with help of A.S. and C.S. All authors read and approved the manuscript.

## Additional information

## Competing financial interests

The authors declare no competing financial interests.

## Materials & Correspondence

All data and code used in this manuscript are available from the corresponding author upon reasonable request.

