## Supplementary Information for "Robustness and timing of cellular differentiation through population-based symmetry breaking"

April 26, 2019

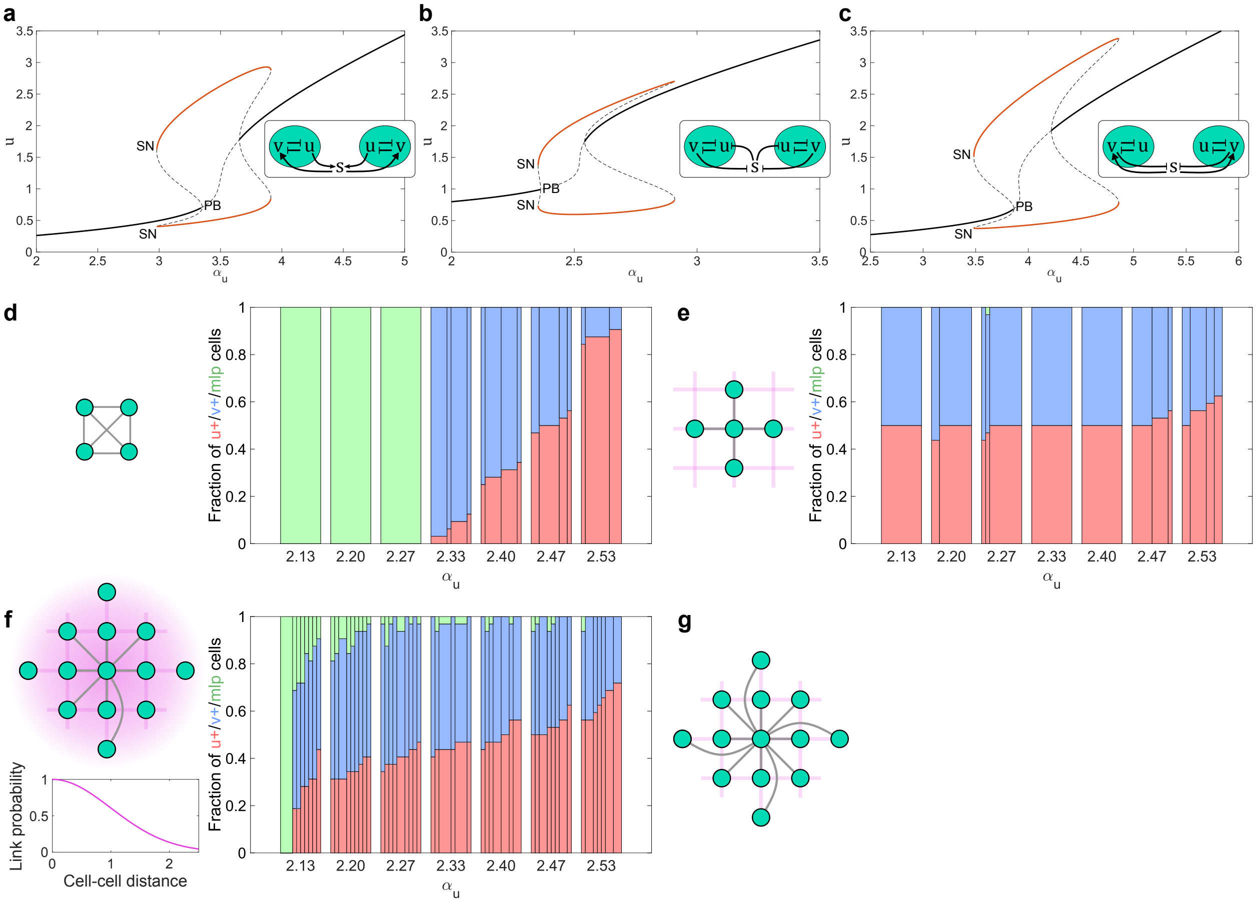


**Supplementary Fig. 1** General features of population-induced symmetry breaking are conserved for different coupling scenarios. **a**, **b**, **c** Symmetry breaking via a population-based pitchfork bifurcation is conserved for different network topologies (schemes in insets and models as described with Eq. (2) in Methods). Bifurcation analyses for two coupled identical cells are shown. PB: symmetry breaking pitchfork bifurcation; SN: saddle-node bifurcation. Solid lines depict stable: homogeneous steady state (HSS, black) and IHSS (red); Dashed lines: unstable steady states. **d**, **e**, **f** u+/v+/mlp cell proportions for increasing $\alpha_u$, when a globally (**d**), locally (**e**) and irregularly coupled (**f**) populations of $N = 32$ cells on a 4×8 grids are considered. The width of each sub-bar within a bar reflects the fraction of occurrence of the respective u+/v+/mlp proportion in the 10 independent realizations. The initial conditions were randomly drawn from a normal distribution $\mathcal{N}(\mu_{ics}, \sigma_{ics}^2)$ around the corresponding $\alpha_u$-specific mlp state as mean ($\mu_{ics}$), and $\sigma_{ics}$=0.1. Model parameters in Methods. **g** Schematic representation of non-local coupling, as used in Fig. 1g.

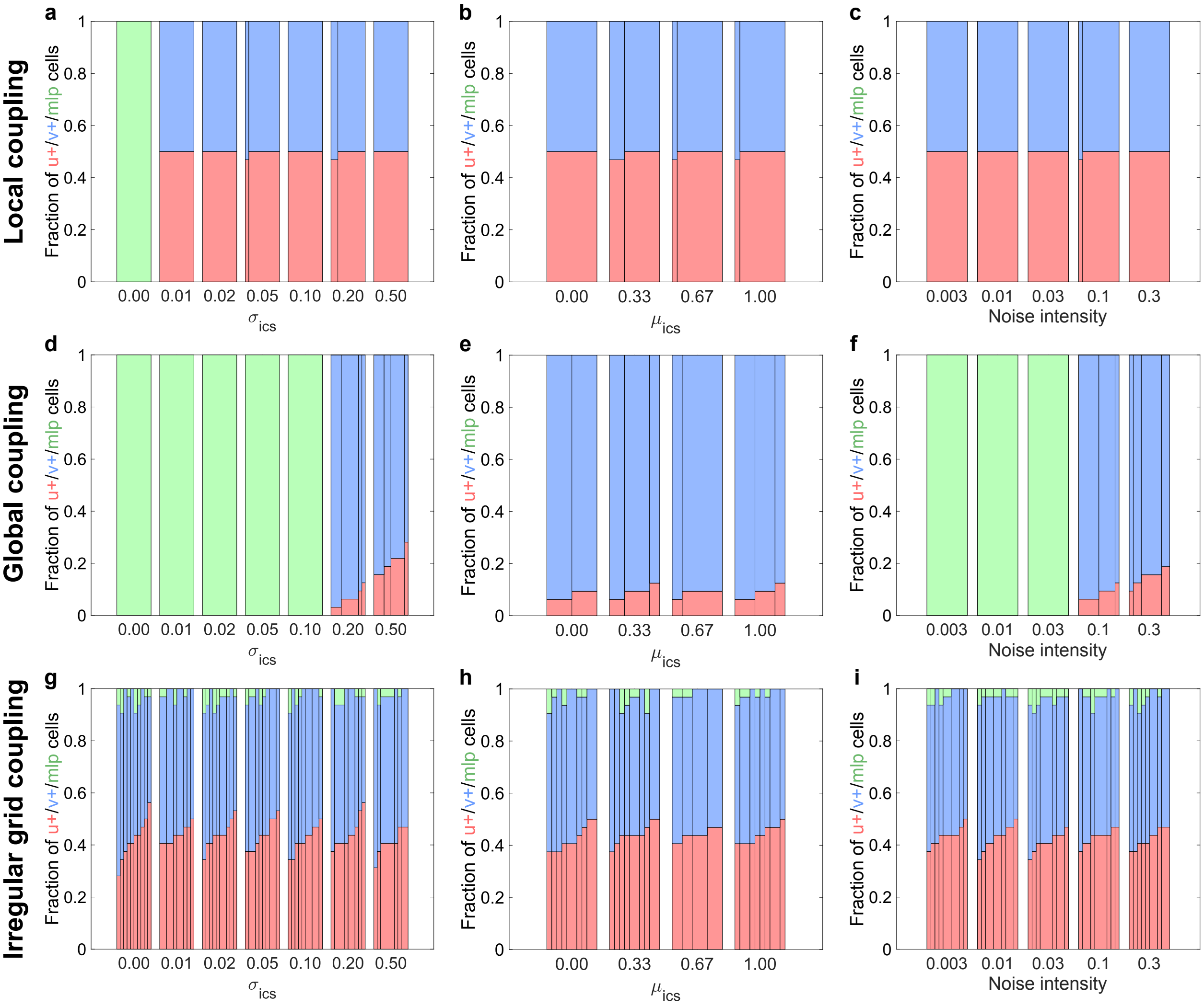


**Supplementary Fig. 2** Robustness in cell ratios is preserved for different coupling schemes. u+/v+/mlp cell type proportions for: **a** Gradual increase in the variance ($\sigma_{ics}$) of the initial conditions mean value around the mlp-state; **b** Shift in the distribution's mean ($\mu_{ics}$) initial value from high v-expression to high u-expression state for stochastic realizations; and **c** Increase in the noise intensity for population of N=32 locally-coupled cells. The bar graphs were calculated as in Fig. 1g, for $\alpha_u = 2.3$. **d**, **e**, **f** u+/v+/mlp cell type proportions for global coupling (conditions as in **a**, **b**, **c**, respectively). **g**, **h**, **i** u+/v+/mlp cell type proportions for irregular coupling (conditions as in **a**, **b**, **c**, respectively).
